# Imaging the mechanisms of anti-CD20 therapy *in vivo* uncovers

**DOI:** 10.1101/2020.05.26.116806

**Authors:** Capucine L. Grandjean, Zacarias Garcia, Fabrice Lemaître, Béatrice Bréart, Philippe Bousso

## Abstract

Anti-CD20 monoclonal antibody (mAb) represents an effective strategy for the treatment of B cell malignancies that may involve complement activity, antibody dependent cellular cytotoxicity (ADCC) and phagocytosis (ADP). While ADP mediated by Kupffer cells is essential to deplete circulating tumors, the relative contribution of each mechanism to the elimination of non-circulating targets has yet to be clarified. Using intravital imaging in a model of MYC-driven B cell lymphoma, we establish here the dominance and limitations of ADP in the bone marrow (BM). We found that tumor cells were stably residing in the BM with little evidence for recirculation. To quantify the contribution of different cytotoxic mechanisms *in situ*, we designed a dual fluorescent reporter to track phagocytosis and apoptosis in real-time. ADP by BM-associated macrophages was the primary mode of tumor elimination but was no longer active after one hour, resulting only in partial depletion. Moreover, macrophage density was strongly reduced in tumor-rich regions. Given their sessile phenotype, macrophages primarily targeted neighboring tumors, resulting in a substantial spatial constraint. Overcoming spatiotemporal bottlenecks in tumor-targeting Ab therapy represents a critical path towards the design of optimized therapies.

**Key points:**

- Functional intravital imaging establishes antibody-dependent phagocytosis as the major mechanism acting at the tumor site during anti-CD20 therapy.
- A transient wave of phagocytosis and a limited macrophage density restrict the efficiency of anti-CD20 anti-tumor activity.

## Introduction

Monoclonal antobodies (mAb) targeting tumor surface antigen represent a powerful strategy for the treatment of several types of cancer. Rituximab, a chimeric IgG1 mAb directed against the CD20 molecule, was the first approved anti-tumor therapeutic mAb and has since improved the prognosis of patients with various B cell malignancies^1,2^. Anti-CD20 therapy is now also widely used to treat several autoimmune diseases^2–6^.

Anti-CD20 mAb acts by depleting normal and malignant B cells and multiple studies have provided clues about its mode of action. *In vitro* studies indicated that anti-CD20 mAb can trigger complement-dependent cytotoxicity (CDC)^7–10^, a mechanism possibly favored by the structure of CD20 clustering^11^ by the mAb. Because antibody-dependent cellular cytotoxicity (ADCC) by NK cells has been readily observed *in vitro*, it has often been considered to be the main mechanism of tumor depletion by therapeutic antibodies^2,12–16^. Notably, antibody-dependent phagocytosis (ADP) by macrophages has also been frequently observed *in vitro* and was reported to be more efficient than ADCC on a per cell basis^17^. Preclinical models have strongly supported the role of FcR-dependent mechanisms for anti-tumor activity^6,18–27^. In addition, depletion of macrophages/monocytes in these models has supported their essential role in mediating mAbs therapeutic activity^18,19,21,24,28,29^. Other studies have pointed towards an important role of neutrophils in therapeutic mAb anti-tumor activity^30–32^. In cancer patients, polymorphisms in Fc receptors have been associated with improved therapeutic response to mAb, further supporting the role of Fc-dependent mechanisms *in vivo*^33–36^.

However, despite two decades of clinical use, the respective contribution of each of these mechanisms to the therapeutic response of anti-CD20 mAb has not been fully understood^2^.

As an additional degree of complexity, the mode of action of therapeutic mAb can also vary with the anatomical site. For example, Ly6C^high^ monocyte-derived cells or resident macrophages were found to be the main effectors during mAb treatment of tumors located in the skin and lung, respectively^24^. Moreover, distinct contributions for Fc receptors and the complement pathway have been observed in the spleen versus the bone marrow (BM) in a model of mAb-mediated cell depletion^29^. Finally, the liver is also an important site for therapeutic activity, as we and others have shown that Kupffer cells are essential to deplete circulating malignant targets in response to mAb^21,22,28^. However, in the context of B cell malignancies, many tumor cells may not have the ability to reach the circulation to be targeted by Kupffer cells, raising the question of putative mechanisms acting in tumor-infiltrated lymphoid organs.

Anti-CD20 therapy also exhibits some limitations. While anti-CD20 mAb has contributed to increase patient survival in distinct types of B cell lymphomas, it is most often not curative^2^. Indeed, a large proportion of patients ultimately relapses and several mechanisms of resistance have been identified ^8,15,37–41^. Preclinical models have shown in particular that the BM is a niche for low therapeutic IgG activity^29,42,43^. Strategies to circumvent these limitations included Ab glycoengineering^22,44–48^, combination with chemotherapy^43^ or with phagocytosis checkpoints^49,50^ including anti-CD47 or anti-CD24 antibodies ^30,51–53^.

Together, these observations highlight the importance of identifying the bottlenecks in anti-CD20 Ab therapy. Here, we combined intravital imaging and a fluorescent dual reporter for phagocytosis and apoptosis to characterize the mechanisms and dynamics of anti-CD20 mAb anti-tumor activity in the BM of a B cell lymphoma-bearing mice. We found that ADP by BM-associated macrophages plays a dominant role in tumor cell elimination. Importantly, both temporal and spatial constraints limited the extent of ADP, raising important considerations for treatment optimization.

## Methods

### Mice and cell lines

6-8-week-old C57BL/6J mice were purchased from ENVIGO. Eμ-myc transgenic mice developing a spontaneous Burkitt-like lymphoma^54^ were bred in our animal facility. A Burkitt-like lymphoma B cell line was isolated from male Eμ-myc mice and transduced to express a FRET-based reporter [CFP^(DEVD)^YFP probe (Eµ-myc DEVD cells) or CFP^(DEVG)^YFP probe (Eµ-myc DEVG cells)] and express CD20. These cells were then established in C57BL/6 mice by injecting 1×10^6^ Eμ-myc cells intravenously (i.v.). Mice were examined every day and sacrificed in case of prostration, tousled-hair, weakness, ectopic or nodal tumor mass >1cm or a weight loss >10%. All animal studies were approved by the Pasteur Institute Safety Committee in accordance with French and European guidelines (CETEA 2017-0038). Cells were cultured in RPMI supplemented with 10% heat-inactivated fetal bovine serum, 50 U.mL^-1^ penicillin, 50 μg.mL^-1^ streptomycin, 1 mM sodium pyruvate, 10 mM HEPES and 50 μM 2-mercaptoethanol, and maintained at 37°C and 5% CO2. Cell lines were routinely tested for the absence of Mycoplasma contamination (Venor-GeM Advance mycoplasma detection kit, Minerva Biolabs).

### Treatments

Anti-CD20 mAb (Clone 5D2, mouse IgG2a, from Genentech) was used at 20 μg.mL^-1^ *in vitro.* Mice were injected i.v. with 50µg of 5D2 or 50µg of the isotype control HY1.2 (mIgG2a) when indicated. In some experiments, cells were pre-treated for 1 hour with bafilomycin A1 (Fischer Scientific) at 50 mM before anti-CD20 mAb addition and bafilomycin A1 was left throughout the assay. Tumors were treated overnight with staurosporine (SIGMA) at a final concentration of 1 µM to induce apoptosis.

### Flow cytometry

For flow cytometry analysis, cell suspensions were Fc-blocked using anti-CD16/32 mAb (BioLegend, clone 93) combined to murine serum 4% (Thermo Fischer). Stainings were performed with the following mAb: CD45.2-BUV737 (Clone 104, BD), Ly6G-BUV395 (Clone 1A8, BD), Ly6C-BV785 (Clone HK1.4, BioLegend), CD11b-PerCp5.5 (Clone M1/70, BioLegend), F4/80-AlexaFluor594 (BM8, BioLegend), F4/80-APC (BM8, BioLegend), F4/80-APCCy7 (BM8, Biolegend), mCD20-Alexa647 (Clone SA275A11, BioLegend). Analyses were performed with a LSR Fortessa II cytometer (BD Biosciences) or a Cytoflex LX (Beckman Coulter) and analyzed with FlowJo software version 10.6.2 (BD Biosciences).

### Bone marrow extraction, BMDM generation and ADCP assay

For *ex vivo* analyses, femurs and tibias were isolated from Eµ-myc tumor-bearing mice or control C57BL/6 mice. Bone marrow cells were extracted by centrifugation of the bones into phosphate-buffered saline (PBS) supplemented with 1 mM ethylenediaminetetraacetic acid (EDTA) and 0.5% fetal bovine serum (FACS buffer), and single-cell suspensions were prepared by filtering the cells through 70-µm cell strainers for further experiment. To generate bone-marrow derived macrophages, 20×10^6^ cells were resuspended in RPMI 20% M-CSF (from L929 supernatant) and plated onto a low-adhering 150mmx25 dish treated by vacuum Gas Plasma plate (Corning REF353025) for a total of 7 days. At day 4, supernatant was removed and fresh RPMI 20% L929 was added. Macrophages were then detached by PBS 5mM EDTA for 10 min at 4°C before being plated at 1×10^6^ cells in 12 well plates and left overnight before doing an ADP assay the following day: 1×10^6^ tumor cells were added to the well and left 30 min at 37°C to settle down before being treated with indicated treatment for 2 hours at 37°C. Phagocytosed tumor cells were quantified by flow cytometry.

### Intravital two-photon imaging

Bone marrow intravital imaging was performed on tumor-bearing mice between 3 to 4 weeks following tumor injection. Briefly, mice were anesthetized with a mixture of Xylazine (Rompun®, 10 mg/kg) and Ketamine (Imalgene®, 100 mg/kg), which was replenished hourly. The scalp hair was removed, and the skin was incised at the midline to expose the bone. The jaw was fixed on the surface of a steel plate to maintain the superior part of the skull horizontally, and a round 20 mm coverslip was centered and fixed above the frontoparietal suture after PBS deposition, using a cyanoacrylate-based glue. During imaging, mice were supplied with oxygen and their temperature was maintained at 37°C with a heating pad. Two-photon imaging was performed with an upright microscope FVMPE-RS (OLYMPUS) and a 25x/1.05 NA water-dipping objective (OLYMPUS). Excitation was provided by an Insight deep see dual laser (Spectra physics) tuned at 820 nm. To create time-lapse sequences, we typically scanned a 30 to 50 μm-thick volume of tissue at 5 μm Z-steps and 75 seconds intervals. The following filters were used for fluorescence detection: CFP (483/32), YFP (542/27) and Alexa 594 (593/35). Movies were processed and analyzed with Fiji software (ImageJ 2.0). Movies and figures based on two-photon microscopy are shown as 2D maximum intensity projections of 3D data.

### Fixed tissue section of the bone marrow and confocal microscopy

Tibia were dissected and left for 4 hours in paraformaldehyde 1% and subsequently transferred to EDTA 0.35M for decalcification. Tissues were dehydrated in sucrose 20% and frozen in OCT embedding compound (Tissue-Tek, Sakura Finetek). 8-µm-thick tissue sections were rehydrated and Fc-blocked with normal mouse and rat serums in the presence of 0.3% Triton X100 (Sigma-Aldrich). Tissues were stained with F4/80-APC (Clone BM8, BioLegend) and tumors were visualized using their CFP or YFP fluorescence. Sections were imaged using confocal microscope Leica TCS SP5. Images were processed using Fiji software (ImageJ 2.0).

### Statistical analysis

All statistical tests were performed with Prism v.8 (GraphPad). A Log-rank, Mann-Whitney, or one-way ANOVA were used as indicated. Data are expressed as mean ± SEM.

## Results

### B cell tumors reside stably in the bone marrow

To study the dynamics of anti-CD20 therapy in a tumor setting, we relied on a mouse model of Burkitt-like B cell lymphoma. We isolated B cell tumors from Eµ-myc mice^54^ that develop spontaneous B cell lymphomas. Tumors isolated from distinct animals were found to express variable levels of CD20 (**Figure S1**). To ensure that CD20 expression was not limiting in our model, we generated by retroviral transduction a B cell tumor cell line expressing high levels of CD20 (in the same range as endogenous B cells *in vivo*) together with a fluorescent protein (**Figure 1A, Figure S2**). When injected in immunocompetent recipients, tumors established primarily in the bone marrow (**Figure 1B**) and were also detected (often at later times) in other organs^55^. In addition, a fraction of tumor cells was found circulating in the blood (**Figure 1B**). Previous studies have demonstrated that circulating tumors are eliminated by Kupffer cells during mAb therapy^21,22,28^. However, the dynamics and fate of B cell tumors residing in lymphoid organs such as the bone marrow is yet to be fully characterized. To address this question, we used intravital two-photon imaging focusing on the bone marrow and the liver, a site where tumor cell recirculation can be readily observed ^21,28^. As expected, a substantial fraction of tumor cells was found rapidly circulating in the liver sinusoids while others were sparsely distributed in the liver parenchyma (**Figure 1C, Movie S1**). By contrast, tumors grew at high density forming clusters in the bone marrow with the vast majority being sessile and showing little evidence for recirculation (**Figure 1C, Movie S2**). Thus, most lymphoid-resident tumor cells are unlikely to reach the liver for Kupffer cell-mediated depletion, reinforcing the key complementary role of local depletion mechanisms during anti-CD20 therapy.

**Figure 1.**
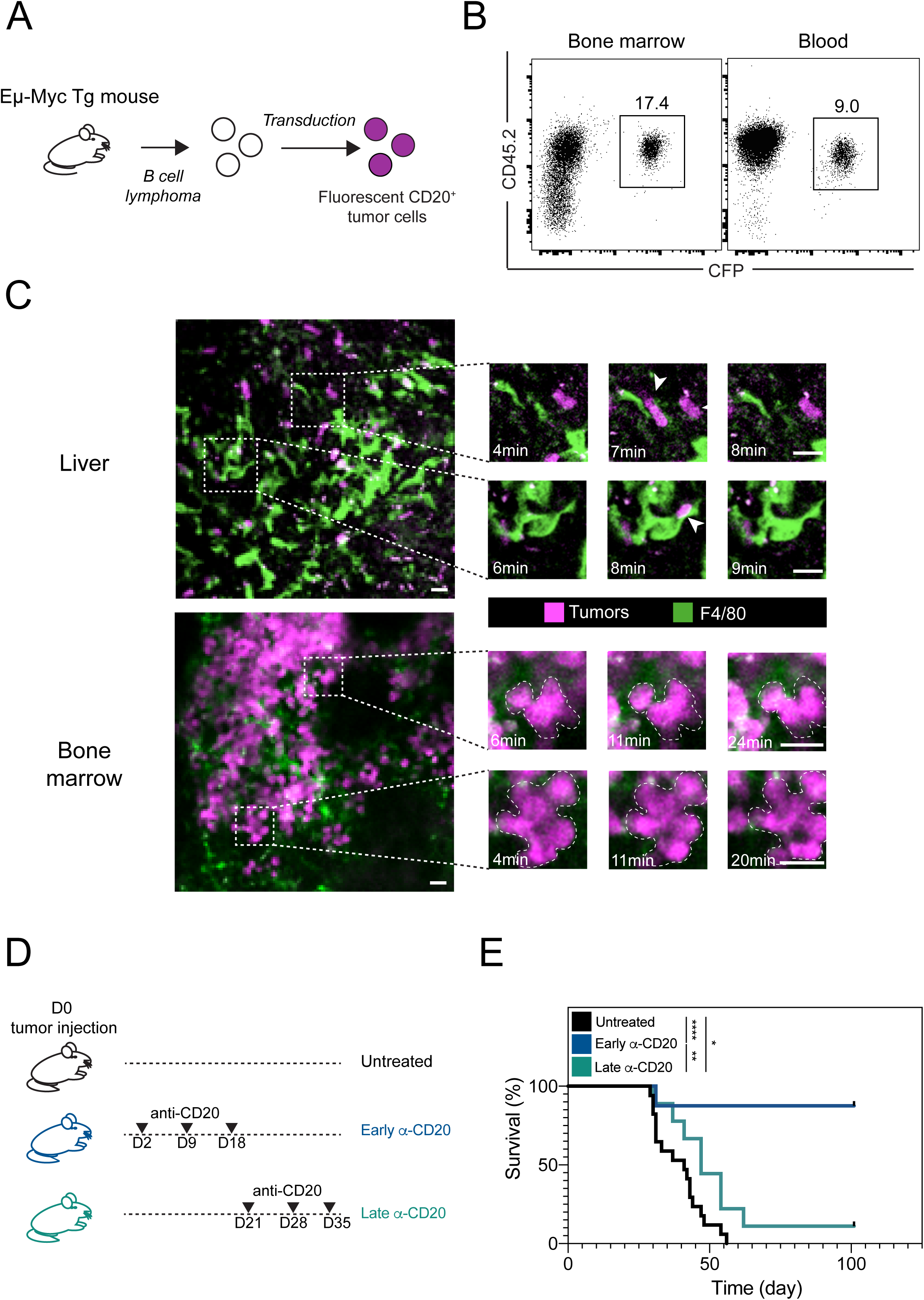
Distinct motility patterns of lymphoma B cells in different anatomical sites. (**A**) Experimental set-up. Eµ-Myc lymphomas were established by intravenous injection of 1×10^6^ fluorescent-CD20^+^ tumor cells in C57BL/6 mice. Recipient mice were analyzed three to four weeks later. Fluorescent CD20-expressing tumors cells were generated by retroviral transduction of B lymphoma cells isolated from Eµ-myc transgenic mice. (**B**) Representative FACS plots showing cells from the bone marrow and the blood of a tumor-bearing mouse gated on CD45.2^+^ cells. Data are representative of 4 independent experiments with n=12 mice analyzed. (**C**) Representative 2-photon time-lapse images of the liver and the bone marrow of tumor-bearing mice. Images illustrate that circulating tumors (white arrows) are detected in the liver sinusoids but that tumor cells stably reside in the bone marrow. Tumor cells appear in magenta and F4/80^+^ macrophages in green. Scale bar, 20µm. Representative of 3 and 5 independent experiments for liver and bone marrow imaging, respectively (**D-E**) Different outcomes for tumor-bearing mice treated early or late with anti-CD20 mAb. (**D**) Schematic of the survival study and the kinetics of treatments. (**E**) Survival curve for mice treated early or late with anti-CD20 Ab or left untreated. Results are compiled from 2 independent experiments with 8 to 17 mice per group. Log-rank test was used for statistical analysis. ***, p<0.001; **, p<0.01; *, p<0.5.

We next tested the consequence of anti-CD20 treatment in our model. In most preclinical studies, anti-CD20 mAb is administrated very early after tumor inoculation. However, the type and efficiency of depletion mechanisms likely vary once the tumor is fully established, a scenario closer to clinical practice. We therefore compared both early and late anti-CD20 treatment in our model (**Figure 1D**). While early treatment cured most animals, late treatment only delayed tumor progression highlighting limitations in the therapy (**Figure 1E**).

### A genetically-encoded reporter to visualize tumor elimination through apoptosis or phagocytosis

To follow tumor cell fate during anti-CD20 therapy *in vivo*, we thought to combine intravital imaging with a genetically-encoded fluorescent probe. Distinct mechanisms can lead to tumor elimination: in particular ADCC results in caspase 3-dependent target cell apoptosis while ADP relies on cell engulfment and degradation^2^. In order to visualize cell degradation during ADP, we thought to take advantage of the differential pH sensitivity of CFP and YFP to detect tumor cells residing in acidic phagosomes after engulfment. Indeed, based on previously described properties of the CFP and YFP proteins^56^, we expected that YFP fluorescence but not CFP fluorescence will be decreasing following phagocytosis (**Figure 2A**). We chose to rely on a FRET-based CFP^(DEVD)^YFP reporter which beside being a tandem CFP-YFP probe, can also be used to monitor caspase 3 activity reflected by DEVD peptide cleavage (**Figure 2A**).

**Figure 2.**
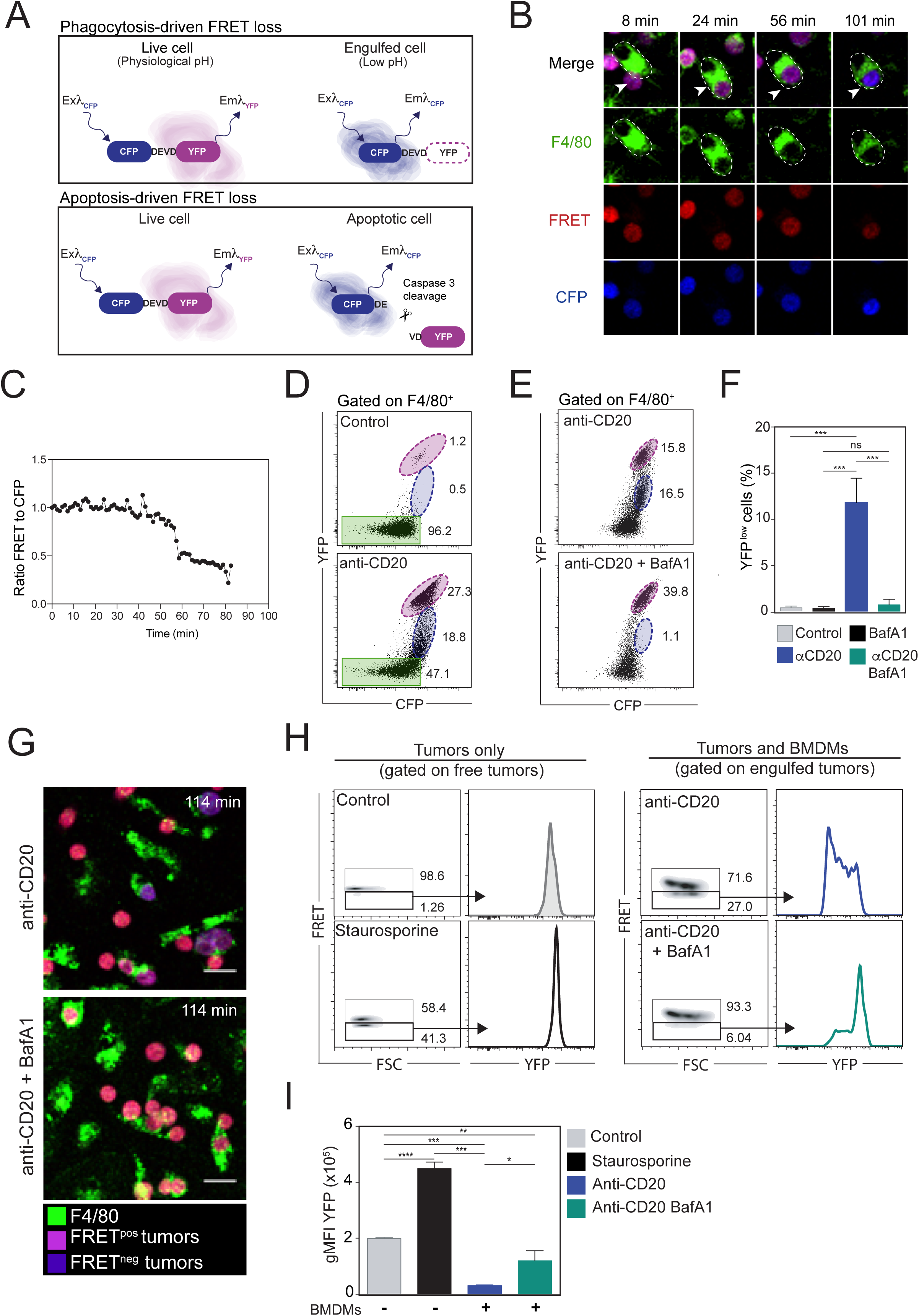
The CFP^(DEVD)^YFP reporter allows real-time visualization of tumor elimination through apoptosis or phagocytosis. (**A**) Schematic illustrating phagocytosis or apoptosis-driven FRET loss in tumor cells expressing the CFP^(DEVD)^YFP reporter. During phagocytosis, the acidic environment causes a loss of YFP signals in tumor cells resulting in FRET loss upon CFP excitation. FRET loss during apoptosis was due to caspase 3-dependent cleavage of the CFP^(DEVD)^YFP reporter, a phenomenon that preserves YFP fluorescence. (**B-G**) Tumor cells were mixed at a 1:1 ratio with BMDMs *in vitro* and cultured in the presence or absence of anti-CD20 mAb and/or BafylomycinA1. (**B**) Tumor cells gradually loose FRET signals after phagocytosis. Representative time-lapse images obtained *in vitro* following anti-CD20 mAb treatment. F4/80^+^ BMDMs are shown in green, FRET signal is shown in red and CFP signal in blue. Upon FRET loss, tumor cells switch from magenta to blue. Representative of 4 independent experiments. (**C**) Quantification of the ratio of FRET to CFP signals over time shown for the engulfed tumor cell in (**B**). Representative of 4 independent experiments (**D**) Detection of anti-CD20-mediated tumor cell phagocytosis by flow cytometry after 2 hours. Two populations of macrophage-associated cells with high (shown in magenta) and low (shown in blue) levels of YFP are detected in the presence of anti-CD20 Ab. Macrophages without tumors are highlighted in green. Representative of 5 independent experiments. (**E-F**) Loss of YFP fluorescence during phagocytosis depends on phagosome acidification. (**E**) Representative FACS plots and bar graphs (**F**) showing the lack of YFP^low^ cells in macrophages treated with BafylomycinA1. ***, p<0.001; ns, is for non-significant (One-way ANOVA) (**G**) Representative time-lapse images showing the lack of FRET loss in tumor cells engulfed by macrophages treated with BafylomycinA1. Data are representative of two independent experiments. Scale bars, 20µm (**H-I**) Detection of apoptosis using the CFP^(DEVD)^YFP reporter. (**H**) Comparison of FRET and YFP signals in cells dying by apoptosis or eliminated by phagocytosis by flow cytometry. Left panels: tumors cells were incubated with staurosporine to induce apoptosis, resulting in FRET loss but maintenance of YFP fluorescence. Right panels: engulfed tumors displayed FRET loss and disappearance of YFP fluorescence. (**I**) Corresponding bar graph showing YFP geometric mean fluorescence intensity for the indicated conditions. Data are representative of 3 independent experiments. ***, p<0.001; **, p<0.01; *, p<0.5.

To validate the use of tandem CFP-YFP probe to monitor ADP, we cultured bone marrow-derived macrophages with CFP^(DEVD)^YFP expressing lymphoma B cells and followed their fate upon addition of anti-CD20 mAb. As shown in **Figure 2B-C**, anti-CD20 mAb promoted rapid and efficient phagocytosis of tumor cells. Importantly, phagocytosed tumor cells exhibited a progressive loss of YFP but not CFP signals as detected by live microscopy and flow cytometry (**Figure 2C-D, Movie S3**). This was not the case for tumor cells prior to phagocytosis. Loss of YFP was also observed with a version of the tandem fluorescent reporter carrying the linker DEVG insensitive to caspase 3 activity, indicating that these events were not related to apoptosis (**Figure S3**). To firmly establish that loss of YFP fluorescence was due to the acidic phagosome environment, we repeated this experiment in the presence of the V-ATPase inhibitor BafilomycinA1 to block phagosome acidification. In this condition, anti-CD20 treatment was equally effective in triggering tumor engulfment but YFP fluorescence remained largely unaffected (**Figure 2E-G**). Thus, by acting as a pH sensor, the CFP^(DEVD)^YFP reporter provide direct evidence for tumor cell phagocytosis and subsequent degradation.

Notably, FRET loss with the CFP^(DEVD)^YFP reporter can also reflect apoptosis through caspase 3 activation and cleavage of the DEVD motif. As shown in **Figure 2H-I**, tumor B cell apoptosis induced by staurosporine also resulted in FRET loss but by contrast to what was observed during phagocytosis, YFP signals could still be detected upon direct YFP excitation. This was expected as the YFP molecule is cleaved from CFP by caspase 3 but remains intact.

Taken together, our results establish that the CFP^(DEVD)^YFP reporter can be used both as a pH sensor during phagocytosis and as a reporter for caspase 3 activity during apoptosis. Additionally, these two modes of cell death should be easily discriminated from one another. Indeed, FRET loss during phagocytosis is a slow process following macrophage engulfment whereas FRET loss by caspase 3 activation is more rapid^57,58^ and should happen before macrophage clearance of the apoptotic body. In the context of anti-CD20 therapy, this reporter should help delineate both ADP and ADCC events in real-time.

### *In vivo* imaging reveals the dominant role of phagocytosis in the bone marrow during anti-CD20 therapy

To measure the local anti-tumor activity mediated by anti-CD20 mAb in the bone marrow, we therefore relied on intravital imaging of tumor cells expressing the CFP^(DEVD)^YFP reporter. We found that anti-CD20 treatment gradually increased the frequency of FRET^neg^ tumor cells in the imaging field (**Figure 3A-B, Movie S4**). Typically, 10-15% of tumor cells exhibited FRET loss after 2hr of treatment, indicative of anti-tumor activity (**Figure 3B**). No increase in FRET^neg^ cells was detected in time-lapse movies performed before anti-CD20 mAb injection (**Figure 3A-B, Movie S4**) or after injection of an isotype control (**Figure 3C**). To clarify the mode of tumor cell death, we labeled macrophages *in vivo* using an anti-F4/80 antibody. With this strategy, we could visualize tumor phagocytosis events in the bone marrow immediately after anti-CD20 mAb administration (**Figure 4A-B, Movie S5-S7**). We also recorded FRET loss in a subset of phagocytosed tumor cells (**Figure 4A and Movie S5-S7**). Notably, not all phagocytosed tumors showed FRET loss at the end of the *in vivo* imaging session most likely because this process required more than 60 min to be detected (**Figure 2C, Movie S3**). Only very few phagocytosis events were observed in the absence of anti-CD20 Ab (**Figure 4B, Movie S5**).

**Figure 3.**
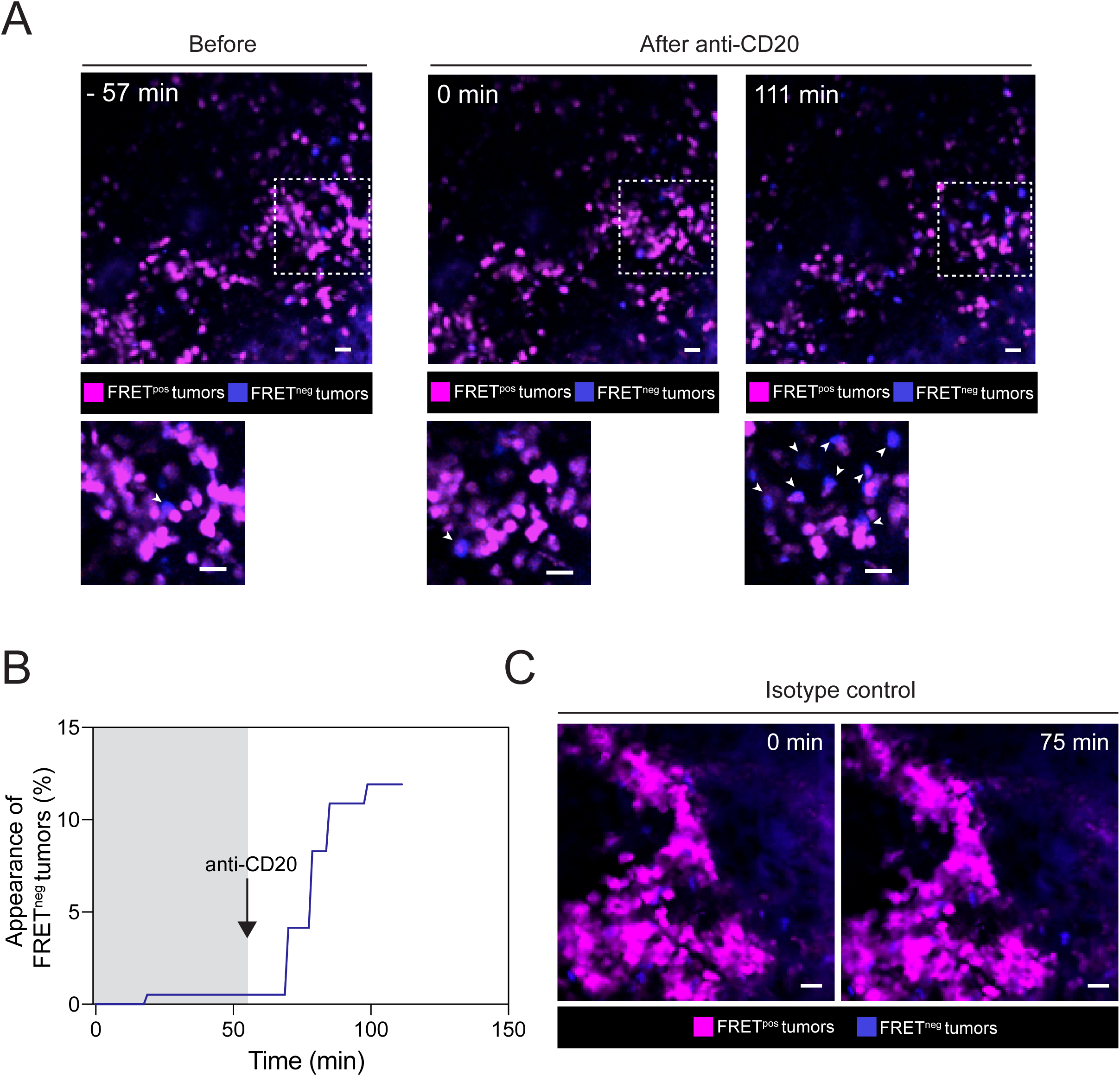
Real-time detection of tumor cell death events *in vivo* following anti-CD20 Ab injection. C57BL/6 mice were injected intravenously with 1×10^6^ tumor cells and were subjected to bone marrow intravital imaging 3 to 4 weeks later before and after anti-CD20 mAb administration. (**A**) Representative two-photon time-lapse images showing the appearance of FRET^neg^ tumor cells (white arrows) following anti-CD20 mAb injection. Scale bar, 20µm. (**B**) Kinetics of appearance of FRET^neg^ tumors before and after anti-CD20 mAb administration as detected by intravital imaging. Data from 3 movies were compiled for quantification. (**C**) Representative two-photon time-lapse images showing that virtually all tumors remain FRET^pos^ in the presence of an isotype control. FRET^pos^ tumors are shown in magenta, FRET^neg^ in blue, Scale bar, 20µm. Data are representative of n>10 movies and from at least 4 independent mice.

**Figure 4.**
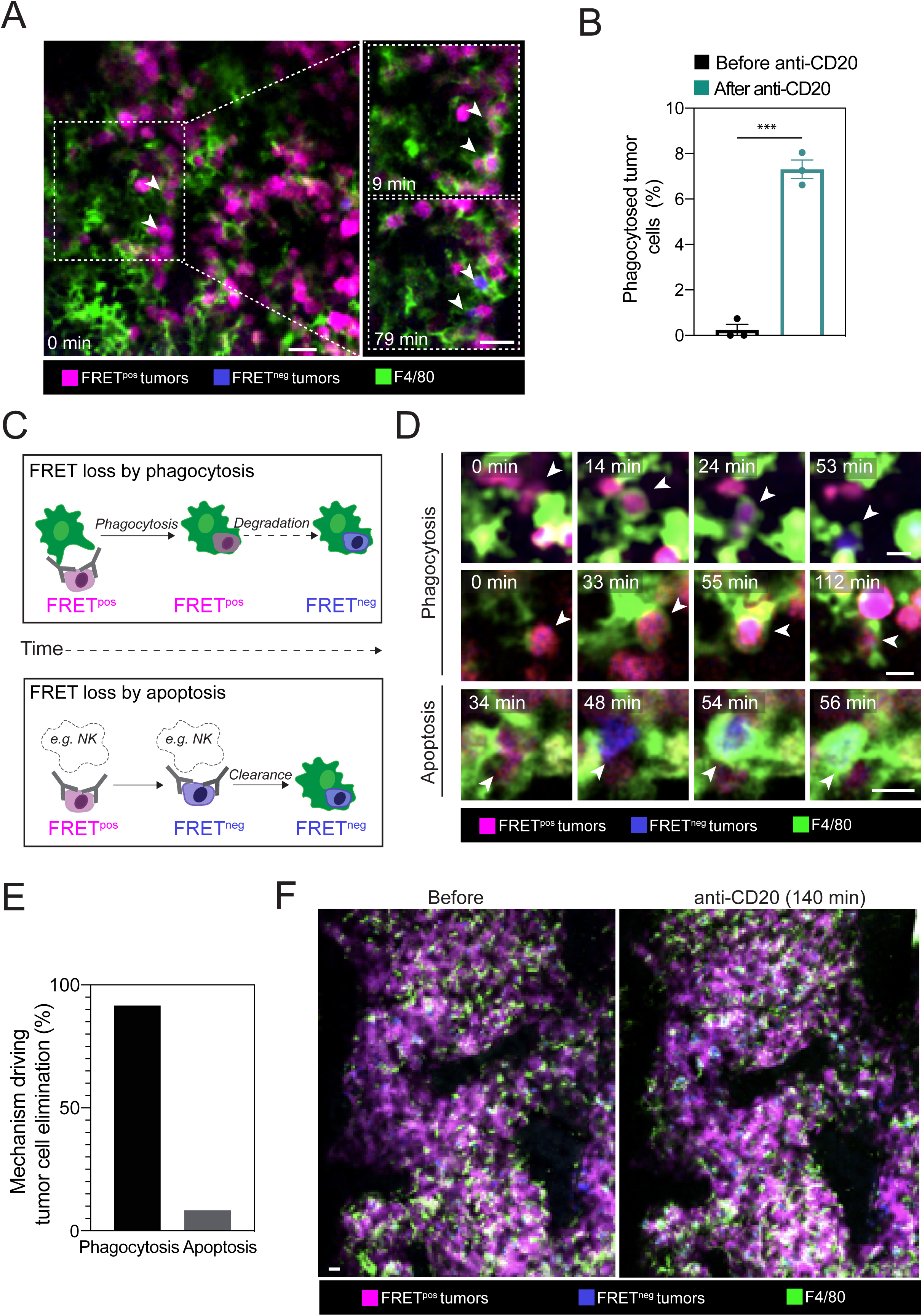
Antibody-dependent phagocytosis is the major mechanism of tumor depletion in the bone marrow in response to anti-CD20 mAb. C57BL/6 mice were injected intravenously with 1×10^6^ tumor cells and were subjected to bone marrow intravital imaging 3 to 4 weeks later just before and after anti-CD20 mAb administration. Macrophages were visualized *in vivo* by injecting a F4/80 fluorescent mAb. (**A**) Representative two-photon images from the bone marrow showing phagocytosis events (white arrow) resulting in FRET loss in engulfed tumors over time. FRET^pos^ tumors are shown in magenta, FRET^neg^ in blue, F4/80-expressing macrophages in green. Scale bar, 20µm. (**B**) Bar graph showing that phagocytosis of FRET^pos^ tumor cells is triggered by anti-CD20 mAb. Phagocytosis events were quantified during a one-hour period either before or immediately after anti-CD20 Ab administration. A unpaired student t-test was carried out for statistical significance. ***, p<0.001 (**C**) Schematic illustrating the different kinetics of FRET loss in tumor cells dying after phagocytosis or apoptosis following anti-CD20 therapy. During ADP, FRET loss is happening after engulfment. Tumor cells dying by apoptosis (due for example to ADCC by NK cells) should lose FRET signals prior to being cleared by a macrophage. (**D**) Representative time-lapse images showing examples of putative ADP or apoptotic events. Scale bars, 20µm. (**E**) Antibody-mediated phagocytosis is the primary mechanism of tumor depletion in the bone marrow following anti-CD20 therapy. Contribution of cell death mechanisms during anti-CD20 therapy. Each cell death event in tumor cells (n= 67) was classified as an ADP or apoptotic event as detailed in (**C**). (**F**) Anti-CD20 therapy shows limited efficacy in the bone marrow. Intravital bone marrow imaging was performed 140 min after anti-CD20 mAb. Note that a large fraction of tumor cells remained FRET^pos^. Scale bars, 20µm. Data in (**A-F**) are representative of n>10 movies and from at least 4 independent mice.

We next thought to more precisely evaluate the contribution of tumor cell death by phagocytosis *versus* apoptosis. We envisioned at least two outcomes (**Figure 4C**). Tumor cells eliminated by phagocytosis should be typically engulfed while being FRET^pos^ and would lose FRET signals only at relatively late time points while located within macrophages. By contrast, tumor cells undergoing apoptosis (e.g triggered by ADCC) should lose FRET signals first and then possibly be cleared by local macrophages. While we found *in vivo* evidence for both outcomes (**Figure 4D, Movie S8**), the vast majority (>90%) of tumor cell elimination could be attributed to ADP, independently of prior apoptosis (**Figure 4E**). We have previously shown that depletion of circulating B cell targets is completed within 2 hours^21,22^. To test whether a similar efficiency is measured in the bone marrow, we examined the fate of tumor cells several hours after anti-CD20 Ab injection. Strikingly, the vast majority of tumors cells were not eliminated at this stage, suggesting some limitations associated with the therapy (**Figure 4F, and Figure S4**). Altogether, these results show that ADP is the main mechanism of action driving tumor cells elimination after anti-CD20 therapy yet may not be sufficient for full tumor eradication.

### Spatiotemporal limitations during anti-CD20 therapy *in vivo*

The partial depletion of tumor cells in the bone marrow was consistent with the moderate improvement in survival seen with late administration of anti-CD20 Ab (**Figure 1A**). To identify potential bottlenecks in anti-CD20 Ab therapy, we first examined the kinetics of ADP following treatment. Compiling 67 phagocytosis events, we observed that tumor engulfment was initiated as early as a few minutes after Ab injection (**Figure 5A**). Importantly, ADP followed a wave-like pattern occurring within a short time frame. ADP events became rare after one hour despite the persistence of macrophages and tumor targets (**Figure 5B**). As shown in **Figure 5C**, cell degradation (as detected by FRET loss) after phagocytosis often required at least an hour *in vivo*. Overall, the transient phase of ADP *in vivo* reveals a temporal limitation of the therapy.

**Figure 5.**
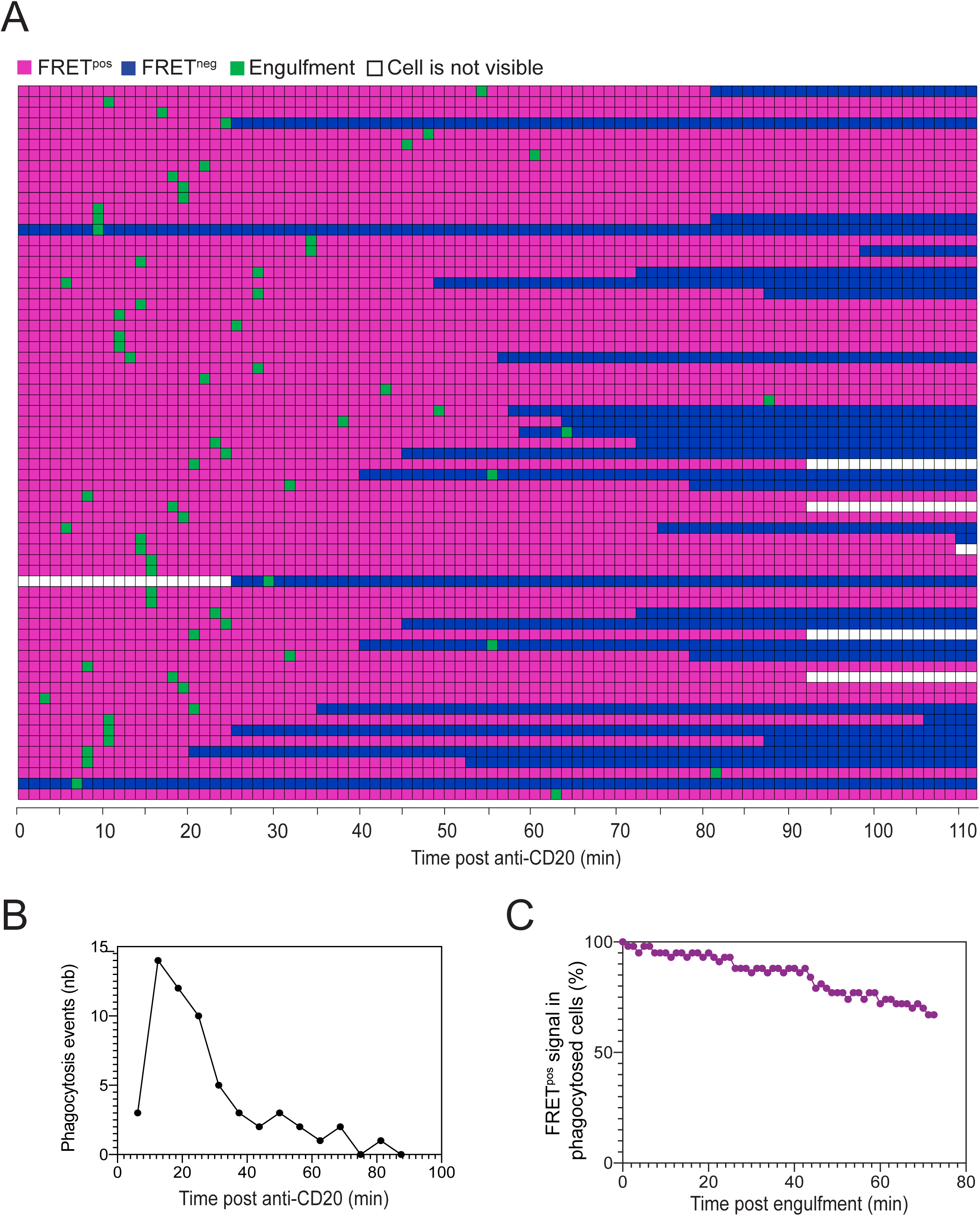
Kinetics of ADP events during anti-CD20 therapy. C57BL/6 mice were injected intravenously with 1×10^6^ tumor cells and were subjected to bone-marrow intravital imaging 3 to 4 weeks later just before and after anti-CD20 mAb administration. (**A**) Kinetics of FRET loss and phagocytosis in tumor cells. Each horizontal line represents a tumor cell eliminated during the imaging period (as detected by FRET loss). Colored squares show the time points during which the tumor is visible within the imaging field. Magenta and blue indicate the time period during which the tumor cell is FRET^pos^ and FRET^neg^, respectively. Green squares indicate the time point at which tumor cells are engulfed by macrophages. (**B**) Kinetics of ADP events after anti-CD20 treatment. (**C**) The timing of FRET loss was quantified for each tumor cell from the time of phagocytosis. Data from (**A-C**) were compiled from n>10 movies obtained from at least 4 independent tumor-bearing mice.

Given the dominance of ADP during response to anti-CD20 mAb, we asked whether the distribution of macrophages within the tumor may also represent a limiting factor for the efficacy of the therapy. Notably, there was a sharp decrease in the density of BM-associated macrophages in tumor bearing mice compared to control animals (**Figure 6A-C**). Specifically, macrophages appeared to be less dense in tumor-rich patches suggesting spatial exclusion (**Figure 6A**). To rule out the possibility that poor macrophage staining *in vivo* was due to limited F4/80 Ab diffusion within dense tumor structures, we performed immunofluorescence on fixed BM-tissue sections. As shown in **Figure 6B** we confirmed that tumor-rich region exhibited a lower macrophage density as compared to non-infiltrated areas of the BM or to tumor-free control animals. Flow cytometric analysis also confirmed that tumor-bearing mice exhibited a decreased percentage of macrophages in the bone marrow (**Figure 6C**). The decreased presence of macrophages likely represents an additional limitation given that BM-associated macrophages appeared largely sessile (**Figure 6D and Movie S9**). To support this idea, we measured for phagocytic events, the distance initially separating the targeted tumor cell from the macrophage center (**Figure 6E**). We found that macrophages could at best reach tumor cells located 30µm away (**Figure 6F**). Overall, our results suggest that macrophage sparseness and low motility together with the sessile behavior of tumor cells may strongly limit the number of tumor cells reachable for ADP. Together, our *in vivo* dynamic analyses uncovered both spatial and temporal bottlenecks in ADP of non-circulating malignant B cells in response to anti-CD20 therapy.

**Figure 6.**
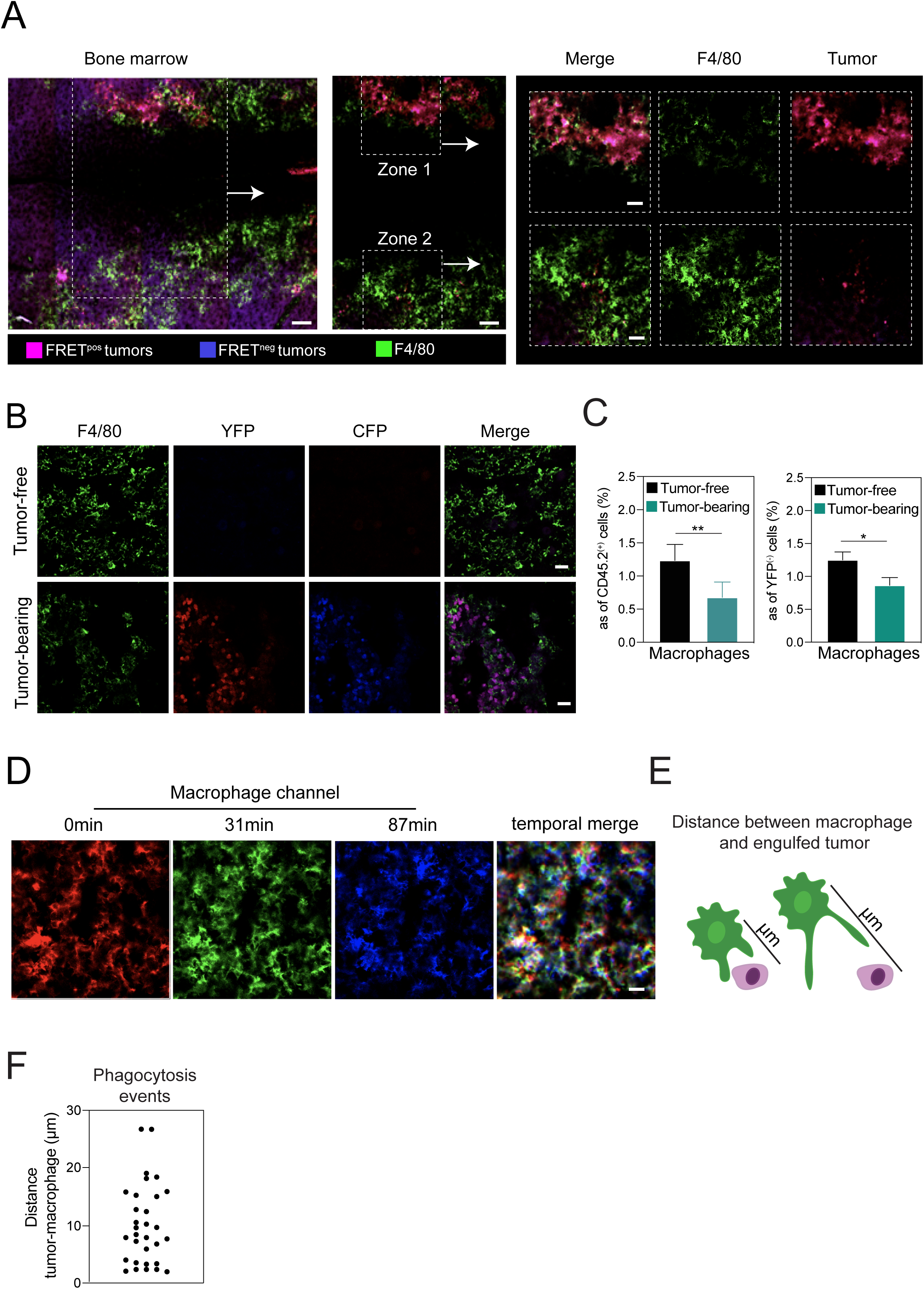
The density of macrophages is a limiting factor during anti-CD20 therapy *in vivo*. (**A**) Representative two-photon image of the bone marrow comparing macrophage density in tumor-rich area (zone 1) and a tumor-free zone (zone 2) within the same imaging field. Scale bar, 100µm; inset scale bar, 50µm. (**B**) The density of bone-marrow associated macrophages in tumor-bearing or tumor-free mouse is shown by immunofluorescence on frozen bone-marrow sections (**C**) Bar graph showing the percentage of macrophages (CD11b^+^Ly6G^-^Ly6C^low^F4/80^+^ cells) in the bone-marrow within total CD45.2^+^ cells or within YFP^-^ cells (excluding tumor cells). Representative of two independent experiments. A Mann-Whitney statistical test was performed for statistical significance. **, p<0.01; *, p<0.5. (**D**) Bone marrow-associated macrophages are mostly sessile. Representative time lapse images of F4/80^+^ macrophages in the bone-marrow. A color was attributed to each time point and overlaid to visualize the migratory phenotype of macrophages. Superimposed signals appear in white and reflect the lack of macrophage motility (**E-F**) Quantification of the distance of macrophage survey. (**E**) Schematic illustrating the distance separating the targeted tumor cell from the macrophage center and (**F**) compilation of distances recorded for individual ADP events.

## Discussion

In the present report, we have established an *in vivo* imaging approach aimed at providing a detailed mechanistic understanding of the anti-tumor activity mediated by anti-CD20 mAb. In a model of MYC-driven B cell lymphoma developing primarily in the bone marrow, we established that ADP but not ADCC was the major mode of tumor elimination. Anti-tumor activity started rapidly after mAb administration but was terminated within 1-2 hours. Moreover, ADP was also limited by the lack of motility of both tumor cells and macrophages. Moreover, this limitation was exacerbated in tumor-rich areas where macrophages were present at low density. Our results support the idea that the rate of ADP is a limiting factor during anti-CD20 therapy.

Several approaches are available to interrogate the mode of action of tumor-targeting mAb therapy. These include *in vitro* experiments with human cells as well as preclinical models relying on cell depletion or genetic ablation of specific receptors (eg. Fc receptors)^18–26^. Intravital imaging provides key additional insights by offering direct visualization of cellular mechanisms and quantitative measurements of cell processes while addressing spatiotemporal variability in therapeutic efficacy^59^. Moreover, the use of functional reporter in imaging experiment can further refine these analyses^60^. While several studies have identified macrophages with engulfed tumors on fixed tissue sections^24,26,43^, it is difficult to formally exclude that tumors were not initially killed by a distinct effector and subsequently cleared by macrophages. Here, we described a fluorescent genetically-encoded reporter to help delineate cell death mechanisms in real-time during mAb therapy. With this probe, we found little evidence of cell death by apoptosis upon anti-CD20 treatment but revealed instead a dominant role for phagocytosis and subsequent cell degradation in tumor elimination. Indeed, of the tumors that were eliminated, more than 90% were targeted by ADP. Thus, our dynamic analyses extend previous studies by providing direct evidence and quantitative measurements for the role of ADP by BM-associated macrophages.

Previous studies have highlighted that anti-CD20 mAb activity can be suboptimal, in particular in the bone marrow^29,42,43^. This activity could be enhanced through chemotherapy^43^ or by interfering with don’t eat me signals^30,50,51,53^. To more precisely identify the origins of these limitations, we established the rate and spatial distribution of ADP events. First, we found that ADP events occurred as a rapid but short wave being terminated 1-2hr after mAb administration. These observations may constitute *in vivo* evidence for saturation of phagocytosis, a phenomenon previously described *in vitro* in which macrophages rapidly lose their phagocytic capacity after an initial round of target engulfment^61^. Thus, strategies aimed at understanding and bypassing this early phase of hyporesponsiveness in ADP may sustain and potentiate anti-CD20 mAb activity. Second, we found that macrophages were largely sessile thus surveying a relatively limited territory for phagocytosis. Indeed, macrophages could target at best tumor cells located at 30 µm from their center. Furthermore, macrophages were found to be less numerous in the bone marrow of tumor-bearing mice and in particular were excluded from tumor rich regions. These findings likely provide a basis for the observed positive correlation between anti-CD20 clinical response and macrophage density in patients with follicular lymphoma ^62^. Strategies to boost the accumulation of phagocytic cells at the tumor site may therefore represent an interesting avenue for improved Ab-mediated cellular toxicity.

In sum, by using functional *in vivo* imaging, we have highlighted the rapid and dominant role of ADP in the bone marrow during anti-CD20 Ab therapy in a model of MYC-driven B cell lymphoma. Furthermore, by uncovering the spatiotemporal dynamics of anti-CD20 anti-tumor activity, we revealed important bottlenecks that should be considered in order to potentiate treatment efficacy.

The ability to measure mAbs functional activity at the single cell level should offer new opportunities to guide the development of novel therapeutic strategies or combination therapies.

## Acknowledgments

We thank members of the Bousso laboratory for critical review of the manuscript. We acknowledge the mouse facility and CB UTechS at Institut Pasteur for support in conducting this study. The work was supported by Institut Pasteur, Inserm and an Advanced grant (ENLIGHTEN) from the European Research Council (P.B).

## Author contributions

C.L.G, Z.G, F.L, B.B conducted the experiments. C.L.G, and P.B designed the experiments, C.L.G. and P.B analyzed the data and wrote the manuscript.

## Conflict of Interest Disclosures

The authors declare no competing interests.

## Notes

### Competing Interest Statement

The authors have declared no competing interest.

