## Supplemental figures and movie legends for "Imaging the mechanisms of anti-CD20 therapy *in vivo* uncovers"

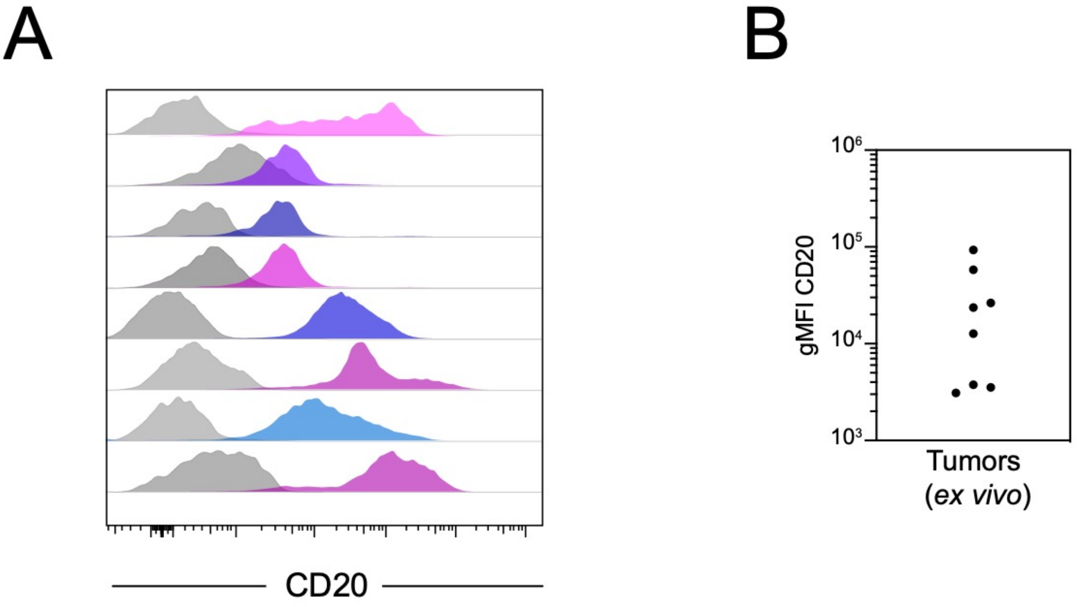


**Figure S1. Spontaneously developing B cell tumors express heterogenous levels of CD20 molecule.** Tumors isolated from Eu-myc transgenic mice were analyzed for CD20 surface expression by flow cytometry. (A) Histograms from multiple individual tumors (magenta to blue) together with unstained controls (gray) are shown. (B) Level of CD20 expression on various Eu-myc tumors are compiled.

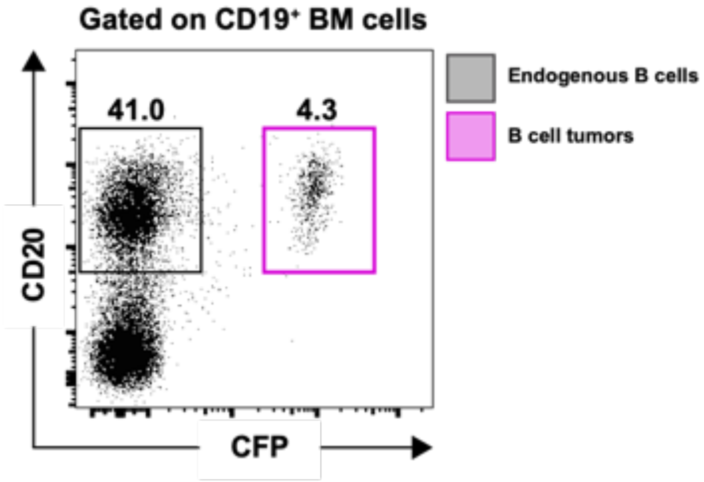


**Figure S2.** ***Ex vivo*** **expression of CD20 on transduced B cell lymphomas.** CFP-expressing Eu-myc tumor cells were transduced to express CD20 and injected in C57BL/6 recipients. After 3-4 weeks, bone marrow cells were analyzed by flow cytometry. The dot plot shows the level of CD20 expression in endogenous B cells and established tumors from the BM.

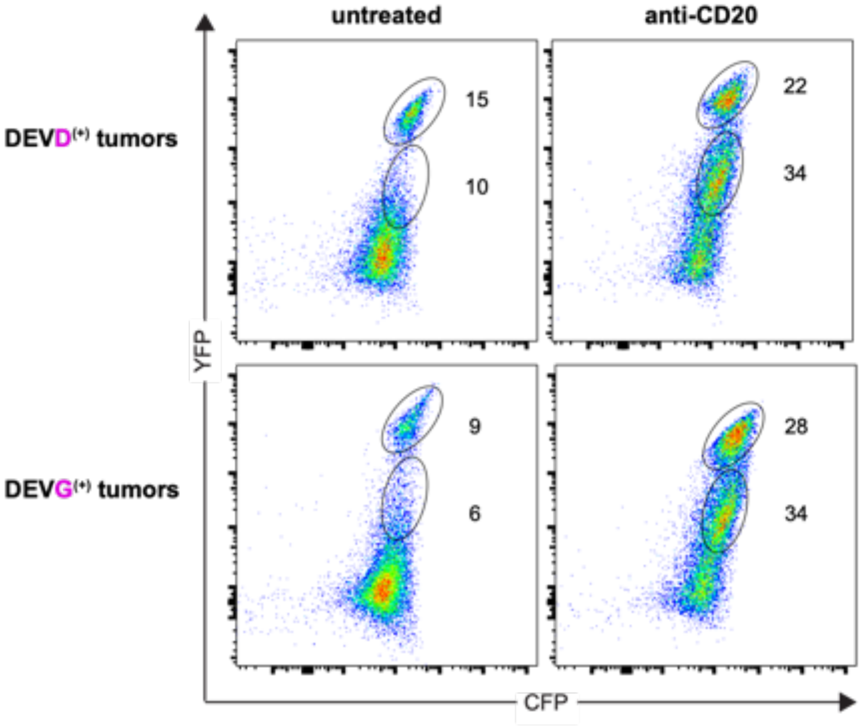


**Figure S3. Loss of YFP fluorescence in phagocytosed tumors is independent of caspase 3 activity.**

Tumor cells expressing the CFP$^{(DEVD)}$YFP or the caspase-3 insensitive CFP$^{(DEVG)}$YFP reporter were mixed at a 1:1 ratio with BMDMs *in vitro* and cultured in the presence (right panel) or absence (left panel) of anti-CD20 mAb. Plots are gated on F4/80-expressing cells. Note that the loss of YFP signal after phagocytosis is not due to caspase 3 activity.

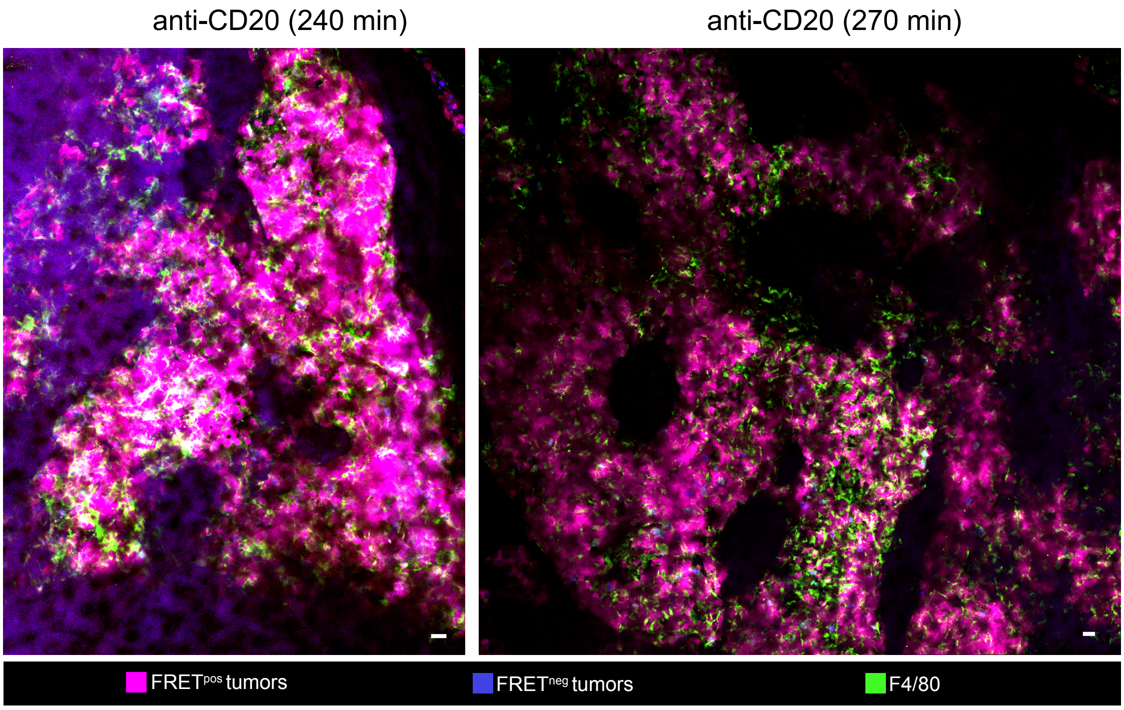


**Figure S4. The vast majority of tumors are not depleted several hours following anti-CD20 treatment.** C57BL/6 mice were injected intravenously with $1x10^6$ tumor cells and were subjected to bone marrow intravital imaging 3 to 4 weeks later just before and after anti-CD20 mAb administration. Anti-CD20 therapy shows limited efficacy in the bone marrow. Intravital bone marrow imaging was performed 240 min or 270 min after anti-CD20 mAbs. Note that a large fraction of tumor cells remained $FRET^{pos}$. Scale bars, 20µm.

### Movie legends

**Movie S1. Detection of tumor cells circulating through the liver sinusoids.**

C57BL/6 mice were injected intravenously with $1x10^6$ tumor cells and were subjected to liver intravital imaging 3 to 4 weeks later before anti-CD20 administration. Tumors are shown in magenta and F4/80$^+$ macrophages in green. Scale bar represent 20µm. Total duration 10 min.

**Movie S2. Bone marrow-associated tumors are mostly sessile.**

C57BL/6 mice were injected intravenously with $1x10^6$ tumor cells and were subjected to BM intravital imaging 3 to 4 weeks later before anti-CD20 administration. Tumors are shown in magenta and F4/80$^+$ macrophages in green. Scale bar represent 20µm. Total duration 31 min.

**Movie S3. A FRET-based reporter to track cell death during phagocytosis in real time.**

Tumors expressing the CFP$^{(DEVD)}$YFP reporter were mixed at 1:1 ratio with BMDMs and imaged in real-time following the addition of anti-CD20 mAb. The movie shows a tumor being phagocytosed and subsequently becoming FRET$^{neg}$. FRET$^{pos}$ tumors are shown in magenta, FRET$^{neg}$ tumors in blue and F4/80$^+$ macrophages in green. Scale bar represent 10µm. Total duration 85 min.

**Movie S4. Real-time detection of tumor cell death events *in vivo* following anti-CD20 Ab injection.**

C57BL/6 mice were injected intravenously with $1x10^6$ tumor cells expressing the CFP$^{(DEVD)}$YFP reporter and were subjected to BM intravital imaging 3 to 4 weeks later before and after anti-CD20 mAb administration. The white circles show the various events of FRET loss detected throughout the imaging time. FRET$^{pos}$ tumors are shown in magenta, FRET$^{neg}$

tumors in blue. The time at which the anti-CD20 Ab is injected is indicated. Scale bar represents 20µm. Total duration 167 min

**Movie S5. Antibody-dependent phagocytosis is the main mode of action of anti-CD20 mAb in the bone marrow: example 1**

C57BL/6 mice were injected intravenously with $1x10^6$ tumor cells and were subjected to BM intravital imaging 3 to 4 weeks later before and after anti-CD20 mAb administration. White circles highlight some events of phagocytosis. $FRET^{pos}$ tumors are shown in magenta, $FRET^{neg}$ tumors in blue and $F4/80^+$ macrophages in green. The time at which the anti-CD20 Ab is injected is indicated. Scale bar represent 20µm. Total duration 167 min

**Movie S6. Antibody-dependent phagocytosis is the main mode of action of anti-CD20 mAb in the bone marrow: example 2**

C57BL/6 mice were injected intravenously with $1x10^6$ tumor cells and were subjected to BM intravital imaging 3 to 4 weeks later immediately after anti-CD20 administration. White circles highlight some events of phagocytosis. $FRET^{pos}$ tumors are shown in magenta, $FRET^{neg}$ tumors in blue and $F4/80^+$ macrophages in green. Total duration 58 min

**Movie S7. Antibody-dependent phagocytosis is the main mode of action of anti-CD20 mAb in the bone marrow: example 3**

C57BL/6 mice were injected intravenously with $1x10^6$ tumor cells and were subjected to BM intravital imaging 3 to 4 weeks later immediately after anti-CD20 mAb administration. $FRET^{pos}$ tumors are shown in magenta, $FRET^{neg}$ tumors in blue and $F4/80^+$ macrophages in green. Scale bar represent 20µm. Total duration 111 min.

**Movie S8. Distinct kinetics of FRET loss *in vivo* were used to distinguished cell death by phagocytosis from cell death by apoptosis**

C57BL/6 mice were injected intravenously with $1x10^6$ tumor cells and were subjected to BM intravital imaging 3 to 4 weeks later before and after anti-CD20 mAb administration. Left panel shows a tumor cell being phagocytosed and subsequently losing FRET signals while the right panel shows a tumor cell losing FRET before being engulfed by a macrophage. $FRET^{pos}$ tumors are shown in magenta, $FRET^{neg}$ tumors in blue and $F4/80^+$ macrophages in green. Total duration 75 min.

**Movie S9. Bone marrow-associated macrophages remain largely sessile *in vivo* after anti-CD20 therapy**

C57BL/6 mice were injected intravenously with $1x10^6$ tumor cells and were subjected to BM intravital imaging 3 to 4 weeks later just after anti-CD20 mAb administration. $F4/80^+$ macrophages are shown in green. Scale bar represents 20µm. Total duration 111 min.
